# Serine catabolism generates NADPH to support hepatic lipogenesis

**DOI:** 10.1101/2021.05.22.445276

**Authors:** Zhaoyue Zhang, Tara TeSlaa, Xincheng Xu, Xianfeng Zeng, Lifeng Yang, Gang Xing, Gregory J Tesz, Michelle F Clasquin, Joshua D Rabinowitz

**Affiliations:** Department of Chemistry, Princeton University, Princeton, New Jersey 08544, USA; Lewis-Sigler Institute for Integrative Genomics, Princeton University, Princeton, New Jersey 08544, USA; Pfizer Inc. Internal Medicine, Cambridge, MA 02139, USA

## Abstract

Carbohydrate can be converted into fat by *de novo* lipogenesis^1,2^. This process is known to occur in adipose and liver, and its activity is upregulated in fatty liver disease^3^. Chemically, *de novo* lipogenesis involves polymerization and reduction of acetyl-CoA, using NADPH as the electron donor^1^. While regulation of the responsible enzymes has been extensively studied, the feedstocks used to generate acetyl-CoA and NADPH remain unclear. Here we show that, while *de novo* lipogenesis in adipose is supported by glucose and its catabolism via the pentose phosphate pathway to make NADPH, liver makes fat without relying on glucose. Instead, liver derives acetyl-CoA from acetate and lactate, and NADPH from folate-mediated serine catabolism. Such NADPH generation involves the cytosolic serine pathway running in liver in the opposite direction observed in most tissues and tumors^4,5^, with NADPH made by the SHMT1-MTHFD1-ALDH1L1 reaction sequence. Thus, specifically in liver, folate metabolism is wired to support cytosolic NADPH production for lipogenesis. More generally, while the same enzymes are involved in fat synthesis in liver and adipose, different substrates are utilized, opening the door to tissue-specific pharmacological interventions.

## Main text

Fatty acids that can be synthesized by *de novo* lipogenesis are called non-essential. Two non-essential fatty acids are highly abundant in mammals: palmitate (C16:0; 16 carbons and zero double bonds) and oleate (C18:1). Together with the essential fatty acid linoleate(C18:2)^6^, these three fatty acids together account for roughly 80% body fat^7^ (Sup. 1a,b).

To quantify their sources in mice fed standard carbohydrate-rich lab chow, we used isotope-labeled water (20% D_2_O)^8–13^. Deuterium from heavy water is stably incorporated into newly synthesized fatty acids both directly and via NADPH^14^. Serum labeling of saponified fatty acids, reflecting both free fatty acids and those covalently embedded in lipids and triglycerides, reached steady state with a half-time of around 1 week (Sup. 1d left), and yet more slowly in white adipose tissue (Sup. 1d right). At steady state, the fraction of each fatty acid coming from synthesis, as opposed to diet, was determined by fitting the observed mass isotopic distribution (Fig. 1a). Labeling of saponified palmitate was extensive across serum and all tissues, while, as expected, linoleate was completely unlabeled (Fig. 1b). Quantitative analysis, summing *de novo* fatty acid amounts across tissues, revealed that most fatty acids in mouse (~80%) come from diet^15^, but a majority of saturated fat is synthesized *de novo* (Fig. 1c). The large contribution of lipogenesis to more atherogenic saturated fat, but not healthier unsaturated fat, may contribute to negative health consequences of high carbohydrate diets and elevated *de novo* lipogenesis^16,17^.

**Figure 1.**
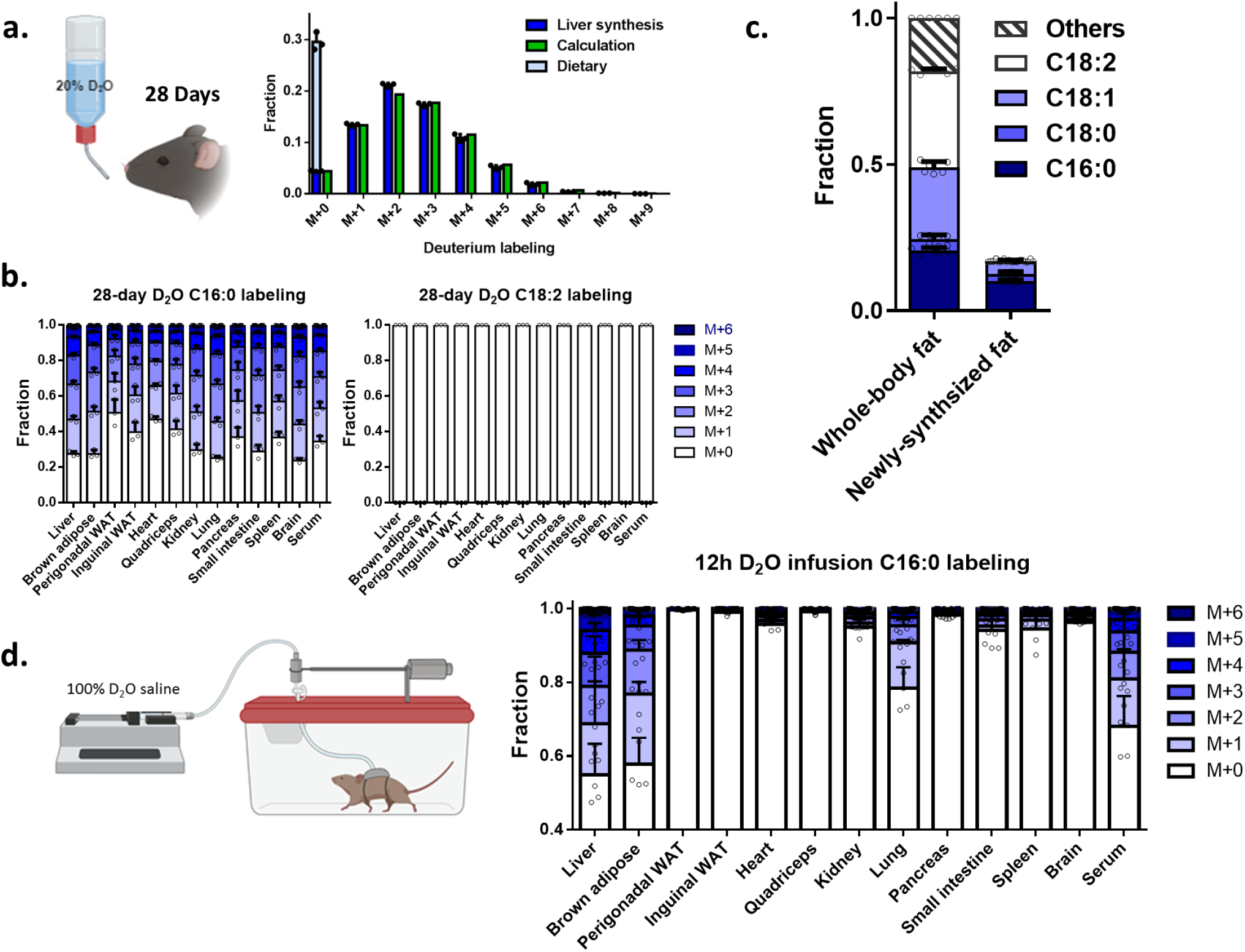
*De novo* lipogenesis in liver and brown adipose produces much of whole-body saturated fat. a. Liver palmitate (C16:0) labeling pattern after 4 weeks of D_2_O drinking. Left bars (blue) reflect the experimentally observed labeling; right bars (green) reflect the calculated labeling based on fraction of newly synthesized fat and observed D_2_O and NADPH labeling. Fraction dietary versus synthesized fat is based on fitting the labeling pattern. Throughout the manuscript, all data are for saponified fatty acids. b. Palmitate (C16:0) and linoleate (C18:2) labeling pattern across tissues after 4 weeks D_2_O drinking. C18:2 is an essential fatty acid and accordingly shows no labeling. c. Composition of whole-body fat (left) and fraction that is endogenously synthesized based on the 4-week D_2_O drinking data (right). d. C16:0 labeling pattern across tissues after 12 h fed-state D_2_O infusion. Mean ± s.d., n=6 mice for composition analysis; n=3 for D_2_O drinking experiment and n=5 for 12h D_2_O infusion.

A challenge in interpreting long term D_2_O labeling experiments is that fat can be exchanged between organs, and indeed the steady-state measurements revealed nearly indistinguishable palmitate labeling across tissues (Fig. 1b, Sup. 1e). To elucidate the organs responsible for lipogenesis, we performed 6 – 12 h infusions of D_2_O^18^. Measurements taken throughout the day at 6 h intervals showed that lipogenesis in mice occurs more rapidly in the night, when mice are more active and eat more (Sup. 1g). Overnight infusions showed greatest saponified fatty acid labeling in liver and brown adipose tissue (Fig. 1d). Only in these two tissues did labeling exceed that of circulating fat (Fig. 1d). This does not rule out the possibility of *de novo* synthesis in other tissues including white adipose, where slow fractional labeling can occur due to the large pre-existing fat pool. Nevertheless, these data show unambiguously that liver and brown adipose tissue actively engage in *de novo* lipogenesis.

Accordingly, we focused on identifying the carbon and hydrogen sources supporting palmitate synthesis in these two tissues. To this end, we infused various ^13^C-labeled nutrients and monitored saponified palmitate labeling (Fig. 2a). We observed a substantial contribution of glucose and lactate to lipogenesis in both liver and brown adipose (Fig. 2b, c; Sup. 2a), whereas acetate contributed strongly only in liver (Fig. 2b; Sup. 2a), indicating an important role of hepatic acetyl-CoA synthetase 2 (ACSS2)^19,20^ which has been proved important in tumor lipogenesis^21,22^.The relative contribution of glucose and lactate differed between liver and brown adipose, with lactate contributing more in liver and glucose more in adipose (Fig. 2b, c, d). Because glucose and lactate interconvert via glycolysis and gluconeogenesis, infusion of either substrate labels both in the circulation^18^ (Fig. 2a; Sup. 2b, c). Accordingly, even if only one of the substrates is used by a tissue to drive lipogenesis, we will observe labeling from infusion of either substrate. By monitoring the extent of tissue palmitate labeling from infusion of each substrate, as well as the extent of cross-labeling between the substrates, the direct contribution of each substrate can be determined by a linear algebra calculation^18^. This analysis revealed that, while glucose and lactate make similar contributions to brown adipose lipogenesis, the contribution of glucose to hepatic lipogenesis is via circulating lactate^18,23^(Fig. 2e). Importantly, there is no discernible direct circulating glucose contribution to liver fat synthesis (Fig. 2e).

**Figure 2.**
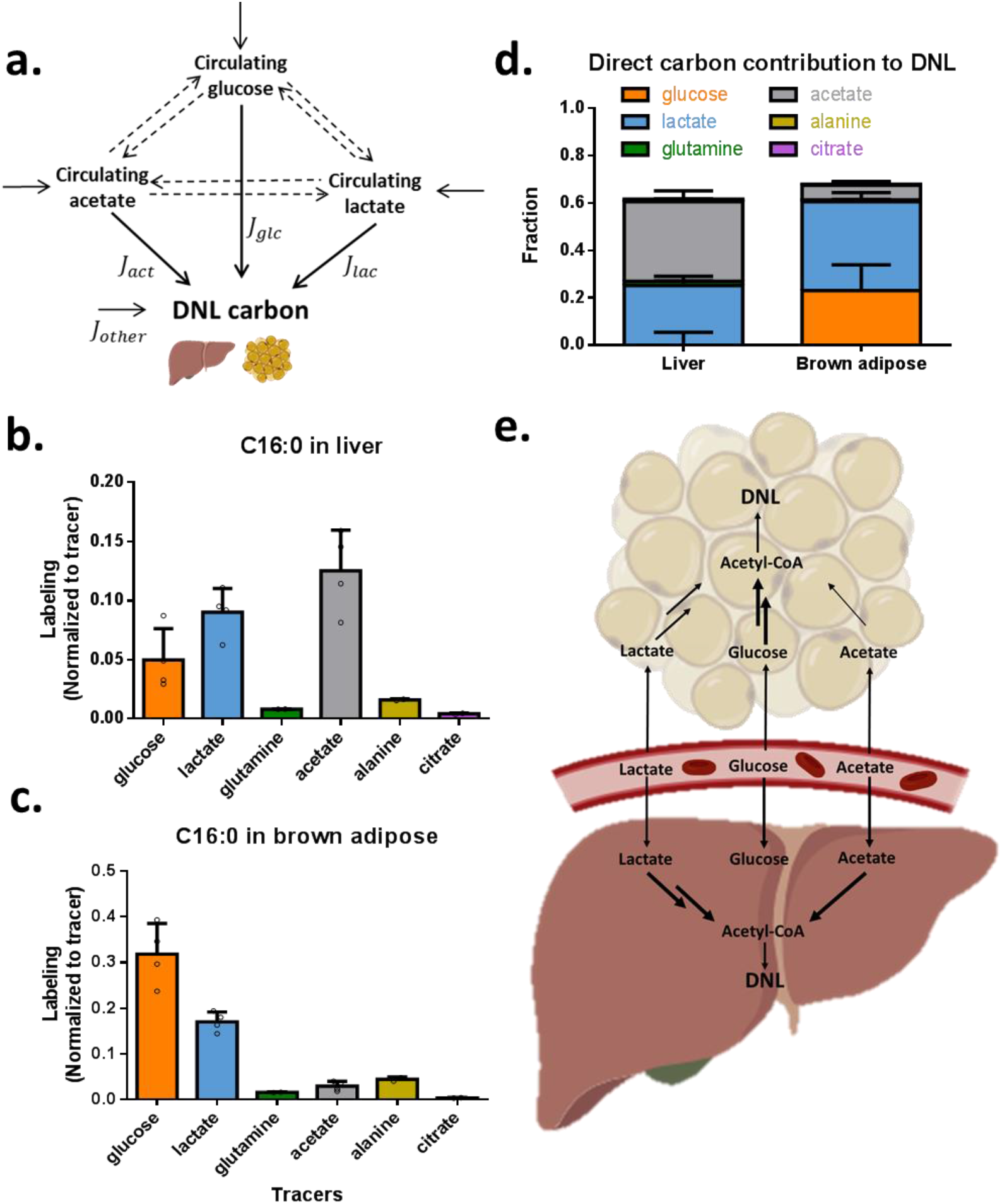
Liver and brown adipose use different carbon sources to support lipogenesis. a. Isotope tracing into fat is complicated by interconversion of the tracer with other circulating metabolites. By conducting tracer experiments with all key circulating substrates, the direct contribution of each can be determined by linear algebra. b. Fractional enrichment of liver C16:0 following 12 h infusion of [U-^13^C]-labeled glucose (n=4), lactate (n=4), glutamine (n=2), acetate (n=4), alanine (n=2) and citrate (n=2). Labeling is normalized to circulating tracer enrichment. c. As in (b), for brown adipose C16:0. d. Carbon sources supporting *de novo* lipogenesis in liver and brown adipose, based on direct contributions to C16:0, calculated based on data in (b) and (c). e. Schematic of differential lipogenic substrate use by liver versus brown adipose. All data are mean ± s.d.

Much of the energy in fatty acids comes from NADPH^1^, which donates hydrogen to drive acetyl reduction. The canonical cytosolic NADPH production route is the oxidative pentose phosphate pathway (oxPPP) ^1,24,25^. Flux from the oxPPP into NADPH’s redox active hydrogen can be traced using glucose deuterated at C1 or C3 (Fig. 3a). Such labeling can be read out in NADPH itself or in downstream products, such as newly synthesized fatty acids. In prior work in cultured cancer cells, we found that NADPH deuterium labeling from the oxPPP is incomplete^26^, but that reflects Flavin enzyme-mediated exchange of NADPH’s active hydrogen with water rather than alternative NADPH sources^14^. Correcting for such exchange, which can be monitored using D_2_O, revealed that the oxPPP is the main cytosolic NADPH source in most cultured cancer cells^2,25,27^. Whether non-transformed lipogenic cells mainly use the oxPPP to generate NAPDH remains unclear.

**Figure 3.**
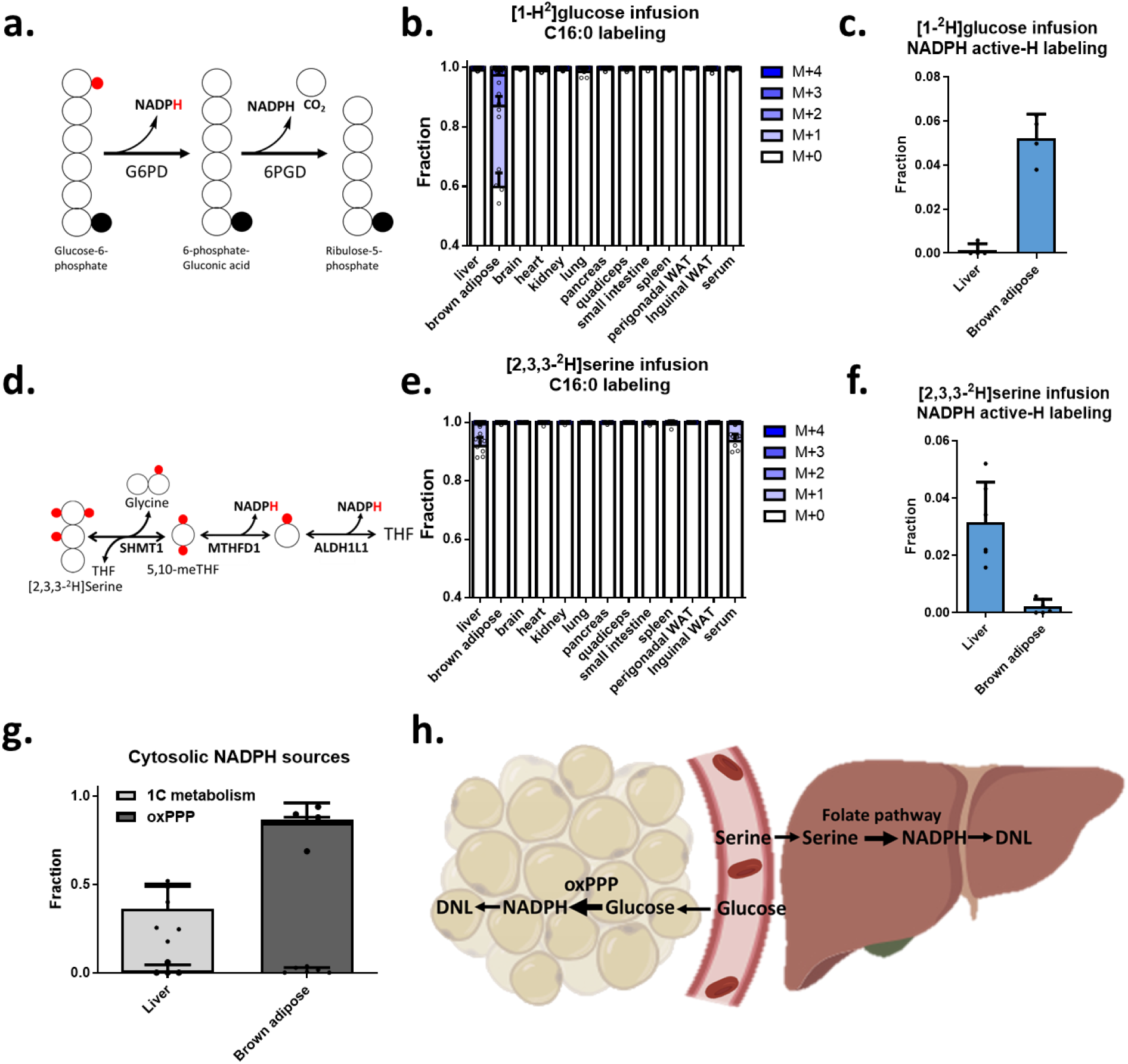
Brown adipose generates lipogenic NADPH via the oxPPP while liver uses serine catabolism. a. Schematic of pathway by which [1-^2^H]glucose and [3-^2^H]glucose labels NADPH via the oxPPP. b. C16:0 labeling across tissues following 12 h [1-^2^H]glucose infusion. c. NADPH active-H measurement in liver and brown adipose from 12 h [1-^2^H]glucose infusion. d. Schematic of pathway by which [2,3,3-^2^H]serine labels NADPH via cytosolic folate metabolism. e. As is (b), for 12 h [2,3,3-^2^H]serine infusion. f. As is (c), for 12 h [2,3,3-^2^H]serine infusion. g. Hydride sources supporting *de novo* lipogenesis in liver and brown adipose, correcting for substrate labeling and H-D exchange between NADPH and water. Calculated based on tracer data in (c) and (f) and D_2_O exchange data in Supplementary Fig.1i. h. Schematic of differential lipogenic substrate use by liver versus brown adipose. Mean ± s.d., n=4 mice for [1-^2^H]glucose infusion; n=7 mice for [2,3,3-^2^H]serine infusion liver, brown adipose and serum; n=4 for other tissues.

To investigate the NADPH sources supporting hepatic lipogenesis, we began by studying cultured primary hepatocytes. Both liver and primary hepatocytes only weakly express oxPPP enzymes^28^. Consistent with this, the absolute oxPPP flux in primary mouse hepatocytes (determined based on radioactive CO_2_ release from [1-^14^C]glucose) (Sup. 3a) was undetectable, despite strong flux in hepG2 cells (Sup. 3b, c). Moreover, in contrast to hepG2 cells, in primary hepatocytes, we observed almost no NADPH and fat labeling from [3-^2^H]glucose, suggesting the predominance of alternative NADPH production pathways^29^ (Sup. 3d, e).

Multiple studies have identified isocitrate dehydrogenase 1 (IDH1) and malic enzyme 1 (ME1) as important cytosolic NADPH producers in lipogenic cells^29,30^. Consistent with this, lysates of primary hepatocytes showed substantial ME1 and especially IDH1 activity (Sup. 3g). In addition, [U-^2^H]glutamine labeled the redox-active hydrogen of NADPH and downstream fat, reflecting cytosolic NADPH production from ME1 and/or IDH1 (Sup. 3d, e). Nevertheless, even after correcting for hydrogen-deuterium exchange based on experiments with D_2_O, the combined contribution of the oxPPP, malic enzyme, and IDH in primary hepatocytes was less than 50% (Sup. 3h).

Serine catabolism is a potential alternative NADPH source, which drives mitochondrial NADPH production for redox defense during cancer metastasis^31^. To date, however, serine catabolism has not been shown to be a physiologically relevant contributor to cytosolic NADPH. We explored whether serine might contribute to NADPH production in primary hepatocytes. Surprisingly, we observed NADPH and downstream fat labeling from [2,3,3-^2^H]serine (Sup. 3d, e).

We attempted to translate the ^2^H-tracing technology *in vivo,* infusing mice with the oxPPP tracer [1-^2^H]glucose overnight. Strikingly, we observed strong labeling of brown adipose palmitate (Fig. 3b), confirming the effectiveness of this tracing strategy. After correcting for the fraction of NADPH’s active hydrogen exchanged with water and the extent of tissue glucose labeling (Fig. 3c), we found that the oxPPP accounts for nearly all brown fat NADPH production (Fig. 3g right). Liver fat was not labeled, however, indicating that liver relies on alternative NADPH sources (Fig. 3b, c).

To identify the source of liver NADPH, we attempted *in vivo* [U-^2^H]glutamine infusion to probe malic enzyme and IDH. Unfortunately, there was extensive deuterium loss between glutamine and malate/isocitrate, rendering the resulting lack of labeling in palmitate relatively uninformative (Sup. 3j, k). The successful tracing with [U-^2^H] glutamine in cultured hepatocytes but not *in vivo* also reflects glutamine being the predominant hepatocytes TCA substrate *in vitro* but not *in vivo^32^* (Sup. 3i). Alternative approaches to label the relevant hydrogen atoms of malate and isocitrate were unsuccessful even in cultured hepatocytes (Sup. 3l).

Accordingly, we turned our attention to serine. Infusion of [2,3,3-^2^H]serine did not detectably label palmitate in brown fat. Impressively, however, we obtained substantial labeling in both NADPH itself and palmitate in liver (Fig. 3e, f).

We next sought to determine the pathway linking serine to NADPH and liver fat synthesis, hypothesizing folate-mediated cytosolic serine catabolism via the SHMT1-MTHFD1-ALDH1L1 reaction sequence^26^. Both MTHFD1 and ALDH1L1 are capable of making NADPH, although neither had been shown to do so in a physiological context previously^33^. Indeed, in most cells, MTHFD1 runs in the NADPH-consuming direction^34^. To assess the direction of one-carbon flux *in vivo,* following [2,3,3-^2^H]serine infusion, we monitored the labeling of 5-methyl-THF. Serine catabolism in mitochondria, followed by re-assimilation of the resulting formate in the cytosol, results in M+1 5-methyl-THF and consumes cytosolic NADPH. In contrast, cytosolic serine catabolism generates M+2 5-methyl-THF and can support cytosolic NADPH production^35^(Fig. 4a). We observed a predominance of M+1 5-methyl-THF in most tissues, but a strong M+2 5-methyl-THF signal specifically in liver, consistent with hepatic one-carbon metabolism producing cytosolic NADPH (Fig. 4b, Sup. 4a).

**Figure 4.**
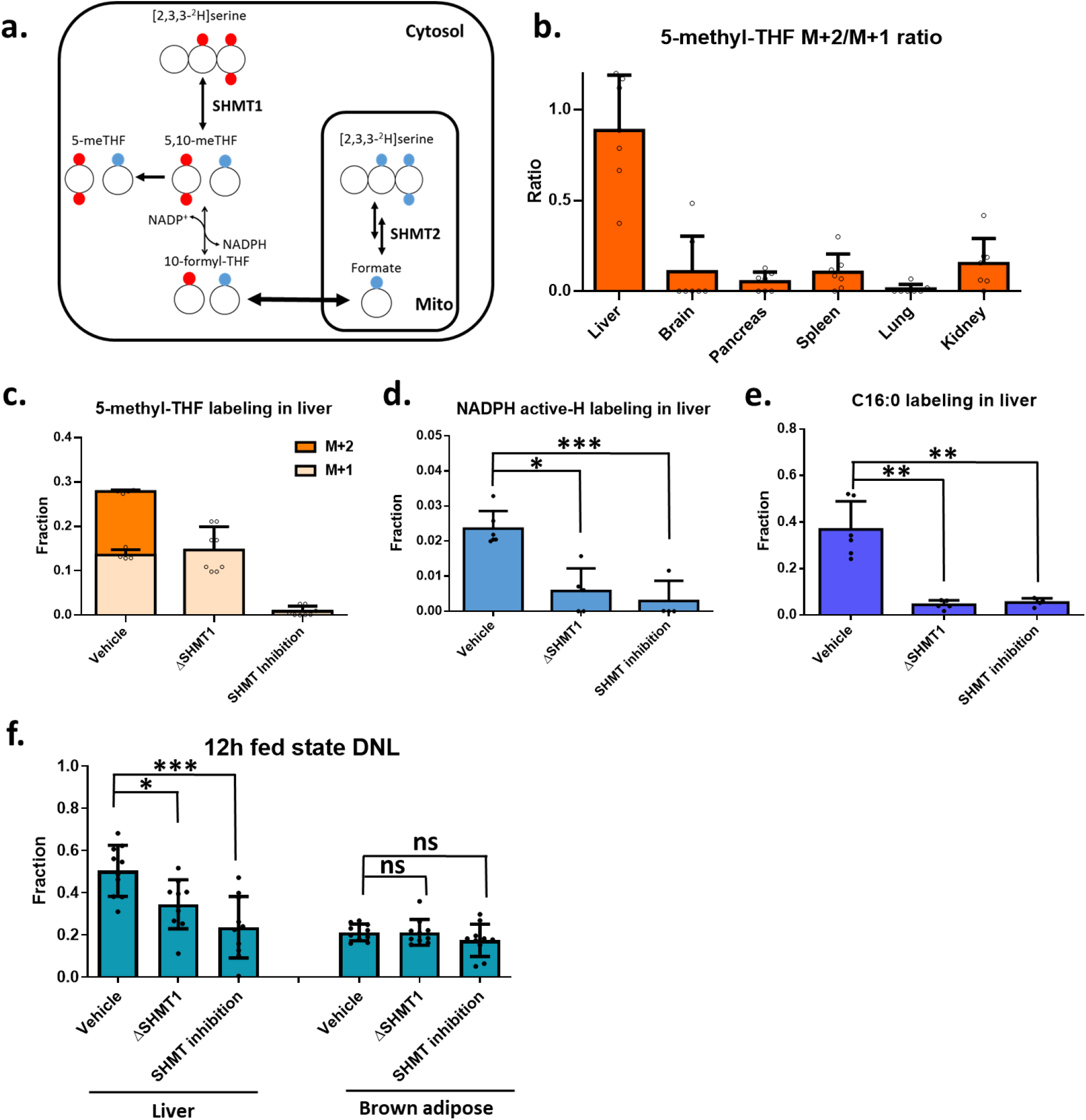
Liver serine catabolism generates NADPH via SHMT1-MTHFD1-ALDH1L1 and thereby supports hepatic lipogenesis. a. Schematic of the cytosolic and mitochondrial serine catabolic pathways. Metabolite labeling from [2,3,3-^2^H]serine generated by the cytosolic pathway is shown in red and mitochondrial pathway in blue. b. 5-methyl-THF labeling for 12 h [2,3,3-^2^H]serine infusion. Ratio of M+2 to M+1 labeling reflects relative contribution of cytosolic versus mitochondrial serine catabolism to 1C units in 5-methyl-THF. c. 5-methyl-THF labeling in liver for 12 h [2,3,3-^2^H]serine infusion in control, whole-body ΔSHMT1 mice, and mice treated with the dual SHMT1/2 inhibitor SHIN2 (3.33 mg/kg/h i.v. infusion). d. NADPH active-H measurement in liver for 12 h [2,3,3-^2^H]serine infusion (vehicle vs ΔSHMT1, p=0.017; vehicle VS SHIN2 treatment, p=0.003) e. C16:0 labeling in liver for 12 h [2,3,3-^2^H]serine infusion. Labeling is normalized to liver serine enrichment and reported as the average number of labeled hydrogens in C16:0. (vehicle VS ΔSHMT1, p=0.004; vehicle vs SHIN2 treatment, p=0.01) f. C16:0 fraction that is newly synthesized in liver and brown adipose overnight, as measured by 12 h D_2_O infusion. (In liver, vehicle VS ΔSHMT1, p=0.028; vehicle VS SHIN2 treatment, p<0.001; in brown adipose, vehicle VS ΔSHMT1, p=0.98; vehicle VS SHIN2 treatment, p=0.19) Mean ± s.d. For serine, N=6 for wild type, N=4 for ΔSHMT1, N=4 for SHIN2 treatment. For D_2_O infusion, N=10 for all 3 conditions. n.s., P>0.05; asterisk, P<0.05; two asterisks, P<0.01; three asterisks, P<0.005.

To confirm that the observed NADPH and fatty acid labeling in liver is derived from cytosolic serine catabolism, we carried out [2,3,3-^2^H]serine tracing in SHMT1 whole-body knock-out mice, which are viable, fertile, and exhibit no major metabolic defects^33,36^. These mice showed a complete loss of M+2 5-methyl-THF (Fig. 4c), confirming functional ablation of SHMT1 enzymatic activity. We also observed an 80% drop in NADPH and palmitate labeling from serine (Fig. 4d, e; Sup. 4c). Thus, SHMT1-driven serine catabolism is a substantial NADPH source for hepatic lipogenesis.

In the absence of SHMT1, folate metabolism can contribute to hepatic NADPH production via ALDH1L1, which can oxidize formyl-THF^37^. Serine can contribute to cytosolic formyl-THF either via SHMT1-MTHFD1 or via the mitochondrial enzymes SHMT2-MTHFD2-MTHFD1L^38–40^. To probe the significance of ALDH1L1, we carried out tracer studies with [^2^H]formate, observing substantial NADPH and fat labeling, although less than that from deuterated serine (Sup. 5c, e). Thus, both the SHMT1-MTHFD1 reaction sequence, and downstream ALDH1L1 which can also be fed via the SHMT2 pathway, contribute to serine-driven hepatic NADPH production.

To more completely inhibit serine’s hepatic NADPH contribution, we used a pharmacological inhibitor of SHMT1/2, SHIN2, a variant of SHIN1 with improved pharmacokinetics^35^. Treatment with SHIN2 markedly decreased both NADPH and palmitate labeling from [2, 3, 3-^2^H]serine in liver (Fig. 4d, e). Labeling from [^2^H]formate, which feeds directly into the formyl-THF pool^37^ and can generate cytosolic NADPH via ALDH1L1 was maintained (Sup. 4e).

Finally, we assessed whether manipulation of serine-driven NADPH production could impact the overall rate of hepatic lipogenesis. D_2_O tracing revealed that both SHMT1 knockout and pharmacological SHMT1/2 inhibition significantly decreased hepatic but not brown adipose fat synthesis (Fig. 4f). Thus, alternative pathways like malic enzyme are insufficient to compensate fully for blockade of serine catabolism, rendering serine catabolism a functionally important driver of hepatic lipogenesis.

One-carbon metabolism is important for organismal development and as a target for cancer therapy and autoimmune disease treatment^4,41–43^. Genetic impairment of mitochondrial serine catabolism (e.g. via knockout of at SHMT2, MTHFD2, or MTHFD1L) results in developmental hallmarks of folate deficiency such as neural tube defects^36^. In cancer cells or T cells, loss of the mitochondrial 1C pathway impairs growth and proliferation^5,31,34,44^. In contrast, SHMT1 loss is tolerated in both development and proliferating cells, rendering the physiological function of cytosolic serine catabolism a long-standing puzzle^36^. The present data show that, while most tissues generate 1C units from serine via the mitochondrial pathway, the liver uniquely relies on cytosolic serine catabolism. Flux running from serine through SHMT1-MTHFD1-ALDH1L1, net generates glycine, CO_2_, and 2 NADPH which support hepatic lipogenesis.

Fatty acid synthesis is a target of interest for treating cancer^45^ and non-alcoholic fatty liver disease^46^. To date, drug development efforts have focused on the core fatty acid biosynthetic enzymes, acetyl-CoA carboxylase (ACC1) and fatty acid synthase (FAS)^46^, which are required for fat synthesis across tissue types and tumors. ACC1 inhibitors have proven effective in decreasing lipogenesis but cause hypertriglyceridemia in both rodents and humans^47,48^. While the mechanism underlying the hypertriglyceridemia remains unclear, a concern with general lipogenesis inhibition is that excess carbon must ultimately be safety stored.

The physiological location for long-term carbon storage is adipose^49,50^. Accordingly, it is conceptually appealing to selectively block lipogenesis in liver, while retaining adipose lipogenesis. The present data show that acetate is a liver-specific lipogenic carbon source, and serine catabolism is a liver-specific cytosolic NADPH source. The associated dependency of hepatic but not adipose lipogenesis on ACSS2 and SHMT1 offers a potential strategy to inhibit selectively hepatic lipogenesis.

## Methods

### Human adipose tissue samples

De-identified human fat samples were the generous gift of Prof. Raymond Soccio and the Human Metabolic Tissue Bank of the University of Pennsylvania Institute for Diabetes, Obesity, and Metabolism.^51^

### Mouse experiments

Mouse work was approved by the Princeton University Institute Animal Care and Use Committee. Mice were housed under normal light cycle (lights on 8:00-20:00) and fed standard chow diet *ad lib* (PicoLab Rodent 20 5053 laboratory Diet St. Louis, MO). Eight-week-old wild type C57BL/6 mice were purchased from Charles River Laboratories. Mice were allowed at least 5 days of acclimation to the facilities prior to experimentation and were randomly chosen for different experimental conditions. No blinding was implemented. Drinking water experiments started one week after arrival and lasted for 4 weeks, with tail vein blood for circulating fatty acid labeling collected weekly.

Intravenous tracer infusions were done on mice after in-house right jugular vein catheterization. Mice were 10-16 weeks old at the time of isotope infusion measurements. Unless otherwise indicated, infusions were performed during the dark cycle from 20:00 to 8:00 +1 day, during which time the mice are more active, feed more, and synthesize more fat. Tail vein blood was collected prior to sacrifice to measure circulating labeling. Blood samples were kept on ice to coagulate and then centrifuged at 4°C to separate serum. Thereafter, mice were anesthetized with isoflurane, opened, and resected. The portal vein was cut, and portal blood was taken with pipet. Then 12 tissues were collected in the following order: liver, spleen, pancreas, kidney, small intestine, perigonadal white adipose tissue, inguinal white adipose tissue, quadriceps, lung, heart, brown adipose tissue, and brain. All tissues were immediately clamped between liquid nitrogen temperature metal plates. After tissue collection, mice were sacrificed by cervical dislocation.

### SHMT1 knockout mice and pharmacological SHMT inhibition

The whole-body ΔSHMT1 mice were a generous gift from Prof. Patrick J. Stover^36^ and bred at Princeton as heterozygotes. Homozygotes knockout progenies and littermate wild-type control mice were used in experiments. The small molecule SHMT inhibitor SHIN2 was mixed with cyclodextrin. Drug or vehicle control was intravenously infused together with tracers through jugular vein for 12 h (20:00-8:00 +1 day) (40 mg/kg of total drug over 12 h, 3.33mg/kg/h). In tracing experiments, no significant differences were observed among true wild type, littermate control and vehicle control mice.

### Primary hepatocyte collection and cell culture

Primary hepatocytes were isolated from wild type C57BL/6 mice^52^ and cultured in high glucose (4.5g/L) DMEM medium supplied with 1% Pen Strep, 100 nM insulin, 100 nM dexamethasone, and 1% glutamax. Primary hepatocytes were cultured in collagen-coated 6-well plates in 37°C, 5% CO_2_ incubator. HepG2 cells were obtained from ATCC and cultured in DMEM with 10% FBS. For labeling experiments, primary hepatocytes were isolated, cultured overnight, and used the next morning. For isotope labeling experiments, hepG2 cells were changed into the same condition as primary hepatocytes 24 h prior to start of tracing. Medium was aspirated and replaced with otherwise identical pre-warmed medium containing the indicated isotope tracer in place of the corresponding unlabeled nutrient. Duration of labeling was 2.5 h for soluble metabolite measurement and 6 h for fat measurement. To harvest metabolites, media was aspirated and 500 μL −20 °C 40:40:20 methanol: acetonitrile: water with 0.5% formic acid was directly added, then neutralized immediately with 15% NH_4_HCO_3_ solution (2.2% v/v of extraction buffer). Cells were collected with a scraper and transferred together with extraction buffer into a 1.5ml vial on dry ice. Samples were incubated on dry ice for 1-1.5 h then thawed on ice and centrifuged at 4°C, 16,000 x g for 10 min. Supernatant was directly analyzed by LC-MS without any washing steps.

### Enzymatic activity assays

A diaphorase-resazurin-coupled biochemical assay was used to detect G6PD, IDH1, ME1 enzyme activity in cell lysates. Cytosolic proteins were isolated by subcellular protein fractionation kit (Thermo, 78840). 2% v/v of cytosolic lysate was mixed with buffer (50 mM Tris pH = 7.4, 5 mM MgCl_2_, 1 mM resazurin, 0.25 mM NADP^+^, 0.1 U mL^−1^ diaphorase and 0.1 mg mL^−1^ bovine serum albumin). To this mixture, substrate (glucose-6-phosphate for G6PD, isocitrate for IDH1 and malate for ME1) was added to a final concentration of 1 mM to initiate reactions. The kinetics of NADPH production were recorded by relative fluorescent unit measurement using a plate reader. The excitation wavelength was 540 nm and emission wavelength 590 nm.

### Radioactive CO_2_ release from oxPPP pathway

^14^CO_2_ release from [1-^14^C]glucose and [6-^14^C]glucose was used to measure oxPPP flux as previously described^2,26^. Briefly, cells were grown in rubber stopper-sealed tissue culture flasks with DMEM containing 0.5 μCi mL^−1^ [1-^14^C]glucose or [6-^14^C]glucose. 150 μL of 10 M KOH was added to a center well (Kimble Chase) containing a piece of filter paper. After 16 h, cell metabolism was quenched, and CO_2_ was released by injecting 1 mL 3 M acetic acid through the stopper. Everything in the center well was transferred into a scintillation vial for counting. Absolute flux was calculated as previously described^2,26^.

### LC-MS sample preparation

Tissue samples were stored at ≤ −70°C. Tissues were pulverized using a Cryomill (Retsch). 10-20 mg of the resulting powder was weighed into a pre-cooled tube for extraction.

Soluble metabolites extraction was done by adding −20 °C 40:40:20 methanol: acetonitrile: water with 0.5% formic acid to the resulting powder (40 μL solvent/mg tissue) and the samples were first vortexed for 5 s and then neutralized immediately with 15% NH_4_HCO_3_ solution (2.2% v/v of extraction buffer). All samples were then vortexed again for 10 s, incubated at −80 °C for 1 – 1.5 h, thawed on ice and then centrifuged at 4°C, 16,000 x g for 10 min. Supernatant was transferred to LC-MS vials for analysis. The acid and associated neutralization are important for NAD(P)(H) measurement and were omitted for ^13^C-labeled samples where NAD(P)(H) measurement was not the focus.

Saponified fatty acid extraction of tissue was done by adding 1 mL of 0.3 M KOH in 90:10 MeOH/H_2_O to the weighed tissue powder. After transferring the resulting mixture to a 4 mL glass vial, saponification was done at 80°C in water bath for 1 h. Then samples were cooled to room temperature, neutralized with 100 μL pure formic acid and extracted with 1 mL of hexane. The hexane layer was transferred to another glass vial, dried down under N_2_, and resuspended in 1:1 methanol: acetonitrile (100 μL/mg tissue; 40 μL/μL serum) for LC-MS analysis.

Serum samples were collected by centrifuging blood at 16,000 x g at 4 °C for 10 min. Then serum was extracted as above for both water soluble metabolites (final volume: 30 μL 40:40:20 methanol: acetonitrile: water per 1μL serum) and saponified fatty acids (final volume: 40 μL 1:1 methanol: acetonitrile per 1 μL serum).

Acetate labeling in serum was measured with a derivatization method: 5 μL serum was added into 100 μL of a mixture of 12 mM 1-ethyl-3-(3-dimethylaminoproryl)carbodiimide (EDC), 15 mM 3-nitrophenylhydrazine, and pyridine (2% v/v) in methanol, incubated at 4 °C for 1 h, and centrifuged for 10 min at 16,000 x g. Then the mixture was quenched with 0.5 mM beta-mercaptothion and 0.1% formic acid in water and directly analyzed by LC-MS analysis^53^.

Folate species were also measured by a derivatization method^54^. 20 mg of tissue was extracted with 1 mL of 1:1 MeOH:H_2_O with sodium ascorbate (25 mM) and NH_4_OAc (25 mM). Precipitates were pelleted by centrifugation (16,000 × g, 10 min). The supernatants were dried down under N_2_ and resuspended in 450 μL of H_2_O with ascorbic acid (30 mM), dipotassium phosphate (50 mM) and 2-mercaptoethanol (0.5%). 25 μL of charcoal-treated rat serum was added to each sample, and the resulting sample was incubated at 37°C for 2 h. Samples were cleaned by Bond Elut PH columns (Agilent Technologies). Columns were washed with 1 mL of MeOH and conditioned with 1 mL of ascorbic acid buffer (30 mM ascorbic acid, 25 mM NH_4_OAc in water). Samples were adjusted to pH 4 using formic acid and were loaded onto the columns. Each column was then washed with 1 mL of the ascorbic acid buffer. Folate species were eluted with 400 μL of 1:1 MeOH:H_2_O with 2-mercaptoethanol (0.5% v/v) and NH_4_OAc (25 mM). The eluate was dried down under N_2_, resuspended in HPLC H_2_O, centrifuged (16,000 × g, 5 minutes), and analyzed by LC-MS.

### LC-MS method for metabolites measurement

LC−MS analysis for soluble metabolites was achieved on the Q Exactive PLUS hybrid quadrupole-orbitrap mass spectrometer (Thermo Scientific) coupled with hydrophilic interaction chromatography (HILIC)^14^. To perform the LC separation of ^13^C-labeled tissue samples, cultured cell samples and all serum samples, an XBridge BEH Amide column (150 mm × 2.1 mm, 2.5 μM particle size, Waters, Milford, MA) was used with a gradient of solvent A (95%:5% H_2_O: acetonitrile with 20 mM ammonium acetate, 20 mM ammonium hydroxide, pH 9.4), and solvent B (100% acetonitrile). The gradient was 0 min, 85% B; 2 min, 85% B; 3 min, 80% B; 5 min, 80% B; 6 min, 75% B; 7 min, 75% B; 8 min, 70% B; 9 min, 70% B; 10 min, 50% B; 12 min, 50% B; 13 min, 25% B; 16 min, 25% B; 18 min, 0% B; 23 min, 0% B; 24 min, 85% B; 30 min, 85% B. The flow rate was 150 μL/min; the injection volume was 10 μL; the column temperature was 25 °C. MS full scans were in negative ion mode with a resolution of 140,000 at m/z 200 and scan range of 75 – 1000 m/z. The automatic gain control (AGC) target was 1 × 10^6^.

Deuterium labeled tissue samples and cultured cell samples are analyzed by an almost identical method with modified gradient and a targeted NADP(H) scan. The gradient was: 0 min, 85% B; 2 min, 85% B; 3 min, 60% B; 9 min, 60% B; 9.5 min,35% B; 13 min, 5% B; 15.5 min, 5% B; 16 min, 85% B, 20 mins stop run, and the injection volume was 15 μL. Full scans as above were alternated with targeted scans: m/z 640-765 with a resolution of 35,000 at m/z=200 (AGC target 5 × 10^5^).

To analyze serum acetate, we also used the Q Exactive PLUS hybrid quadrupole-orbitrap mass spectrometer. LC separation was on a reversed phase column (Acquity UPLC BEH C18 column, 2.1 mm x 100 mm, 1.7 5 μm particle size, 130 Å pore size; Waters, Milford, MA) using a gradient of solvent A (water), solvent B (methanol): 0 min, 10% B; 1 min, 10% B; 5 min, 30% B; 7 min, 100% B; 11 min, 100% B; 11.5 min, 10% B; 14 min, 10% B. Flow rate was 200 μL/min and column temperature was 60°C with an injection volume of 10 μL^53^. MS scans were in negative ion mode with a resolution of 15,000 at m/z 200 and scan range of 100-300 m/z. The automatic gain control (AGC) target was 1 × 10^6^.

To analyze folates^54^, we again used the Q Exactive PLUS hybrid quadrupole-orbitrap mass spectrometer. LC separation was on a different reversed phase column (Agilent InfinityLab Poroshell 120 Bonus-RP 2.7 μm, 2.1 x 150 mm) with a gradient of solvent A (1% vol of 1M NH_4_OAc and 0.1% vol of glacial acetic acid), solvent B (acetonitrile): 4 min, 80% B; 10 min, 2% B; 6 min, 30% B; 11min, 100% B; 15 min, 100% B; 16min, 2% B; 20min 2% B. The flow rate was 200 μL/min and the column temperature was 25°C with an injection volume of 20 μL. MS scans were in negative ion mode with a resolution of 35,000 at m/z 200 and scan range of 350-1000 m/z. The automatic gain control (AGC) target was 1 × 10^6^.

To analyze fatty acids, we used an Exactive orbitrap mass spectrometer. LC separation was via reversed phase-ion-pairing chromatography on a Luna C8 column (150 × 2.0 mm2, 3 μM particle size, 100 Å pore size; Phenomenex) with a gradient of solvent A (10 mM tributylamine + 15 mM acetic acid in 97:3 H2O:MeOH, pH 4.5), solvent B (MeOH): 0 min 80% B; 10 min, 90% B; 11 min, 99% B; 25 min, 99% B; 26 min, 80% B; 30min, 80% B. The flow rate was 250 μL/min and column temperature 25 °C with an injection volume of 5 μl. The MS scans were in negative ion mode with a resolution of 100,000 at m/z 200 and scan range of 120-600. The AGC target was at high dynamic range.

All data from labeling experiments were analyzed by El-MAVEN^55^ and subjected to natural abundance correction^56^.

### Data analysis

#### 1. Quantification of *de novo* lipogenesis flux with D_2_O

##### 1) Long-term steady-state analysis with oral D_2_O

D_2_O labels lipogenic fat both directly and via NADPH. Accordingly, a double binomial distribution model was used to calculate the palmitate labeling pattern based on experimentally measured D_2_O and NADPH labeling^14^. Fractional D_2_O enrichment *in vivo **p_1_*** was determined based on labeling of soluble metabolites pairs (malate-fumarate; glutamate-ketoglutarate). NADPH active-H labeling ***p_2_*** was based on the NADPH-NADP^+^ pair:

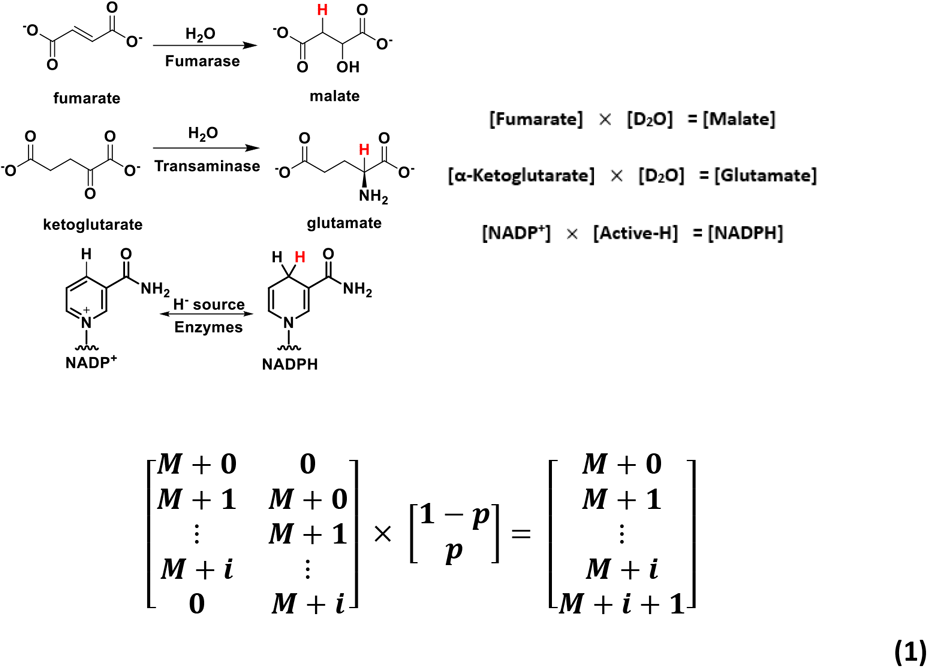

In the above equation, the matrix on the left-hand side contains the experimentally measured mass isotope distribution for the compound on the left-hand side in the equations in the above schematic (the same compound’s mass isotope distribution is in both columns, offset as indicated). The matrix on the right-hand side contains the experimentally measured mass isotope distribution for the compound on the right-hand side in the above equations. The value of ***p*** is determined by solving the resulting linear equations. Values of ***p_1_*** are the average of ***p*** determined using fumarate-malate and α-ketoglutarate-glutamate; ***p_2_*** is solely determined by NADP-NADPH.

For the infusion of other deuterium tracers (1-[^2^H]glucose and [2,3,3-^2^H]-serine), NADPH active-H labeling is also calculated by *eq 1*.

Synthesis of one palmitate molecule (C16:0) requires 7 repeated reactions that incorporate 7 hydrogens from water and 14 hydrogens from NADPH. The expected **(*E*)** number of deuterium in one palmitate is

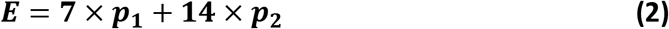

Let ***M_i_*** be the fraction of palmitate containing ***i*** deuterium atoms (i.e. measured fraction at mass *M + i* after correcting for natural isotope abundance). Then ***D*** is the average measured number of deuterium accumulated at steady state per palmitate

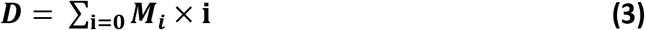

Define:

***F*** = *de novo* lipogenic fatty acid (palmitate) amount at steady state
***C*** = total fatty acid (palmitate) amount
***F/C** =* fraction palmitate synthesized *de novo*

The total deuterium assimilated is given by both (i) total fatty acid amount (**C**) x measured average deuterium per fatty acid (**D**) and (ii) newly synthesized fatty acid amount (to be determined) x expected deuterium per newly synthesized fatty acid (**E**):

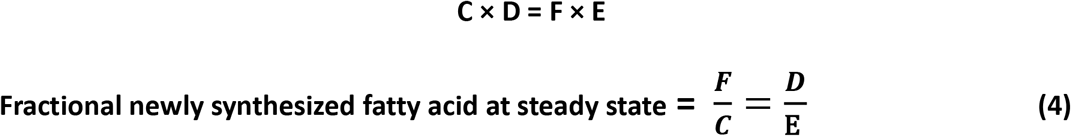

##### 2) Kinetic analysis of *de novo* lipogenesis with D_2_O infusion

For the 12 h infusion experiments, D_2_O in serum is not at steady state and accordingly a slightly more complex calculation is required. D_2_O increases linearly with infusion time (Sup. Fig. 1f). The final serum D_2_O enrichment **(*p_1_*)** can thus be written as a linear function of time:

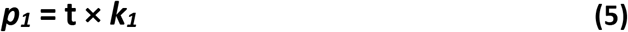

Due to the rapid H-D exchange flux, tissue NADPH labeling reaches equilibrium with serum water labeling quickly, and therefore tissue NADPH will also rise linearly over time:

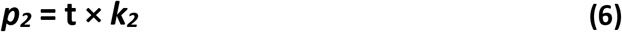

where:

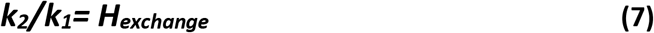

The expected deuterium per palmitate ***(E)*** will be:

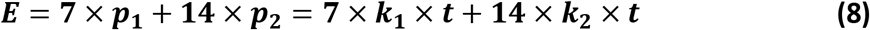

Define *f* (gram/hour) as the average *de novo* lipogenesis flux of palmitate per hour. For an infusion of duration t hours, the total deuterium amount incorporated into newly synthesized palmitate is:

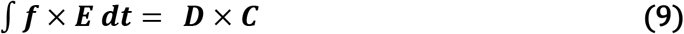

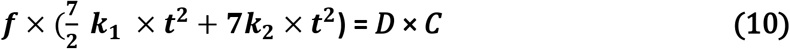

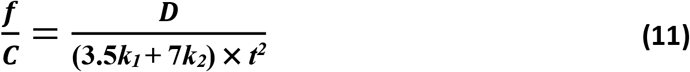

Newly synthesized palmitate ***(F)*** in t = 12 h is given by:

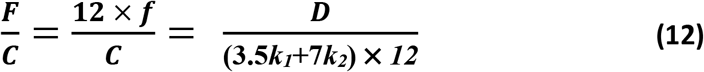

#### 2. Quantification of direct carbon contribution from circulating substrates

The direct carbon sources for lipogenesis can be analyzed similarly to the direct circulating nutrient contribution to TCA^18^. Since *F_circ_^18^* of glucose, lactate and acetate are high, their labeling in serum reaches steady state within ~1 h of infusion. Saponified fatty acid labeling, however, does not reach steady state over 12 h. So, an extra factor of newly synthesized fat (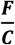 from *eq 12*) needs to be included.

Here, we define ***f_Pal←Met_*** as the carbon flux directly coming from a given circulating metabolite to palmitate. By direct, we mean that the circulating metabolite is taken up into the lipogenic tissue (liver or adipose) and converted within the tissue to acetyl-CoA that is used for fat synthesis. This contrasts with circulating metabolites that are converted into a different circulating lipogenic precursor before entering the lipogenic tissue to support fat synthesis. Here, we assessed six potential lipogenic substrates: glucose (glc), lactate (lac), acetate (act), glutamine (gln), alanine (ala) and citrate (cit).

Define ***L_Met2←Met1_*** as the average pseudo-steady-state ^13^C enrichment fraction per carbon in metabolite_2 after infusion of U-^13^C- labeled metabolite_1 as the tracer. To calculate enrichment, let ***N_i_*** be the fraction of the metabolite containing ***i*** ^13^C atoms (after natural isotope correction) and n the total carbon number of the metabolite:

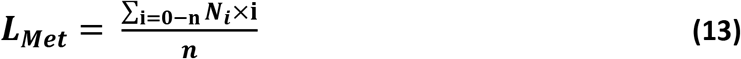

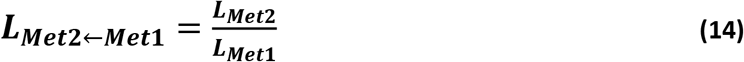

Define the pre-steady-state ^13^C-enrichment measured in saponified palmitate as ***L_pal←Met1_12h_***. The expected pseudo-steady-state palmitate labeling is then given by dividing by the fraction of newly synthesized palmitate over the 12 h:

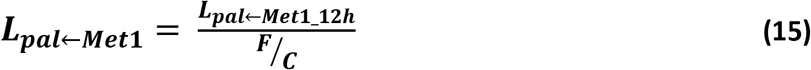

Liver is fed by 22% systemic arterial blood and 78% portal vein blood^57^; brown adipose tissue is fed by arterial blood^53^. Artery enrichment was calculated based on tail vein blood enrichment^18^, and portal vein enrichment was directly measured **(Sup. 2c)**. We can then solve the direct carbon flux to *de novo* lipogenesis in liver and brown adipose tissue with the following equation:

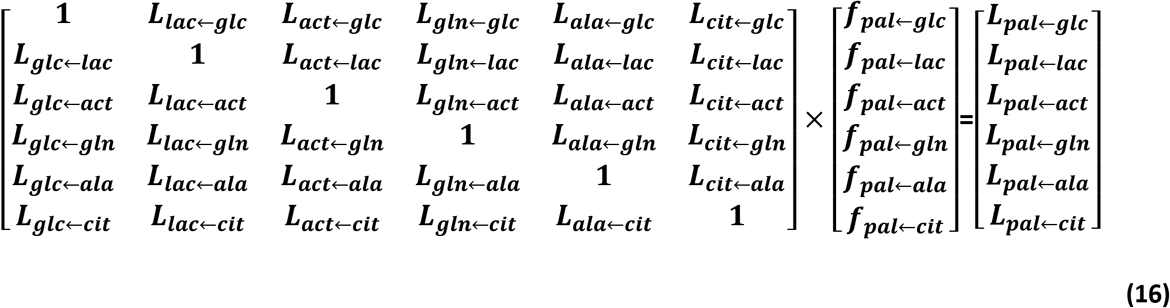

#### 3. Quantification of fractional NADPH contribution from different production pathways

Deuterium-labeled metabolite tracers label NADPH’s active-H^14^. Active-H labeling in NADPH was determined from the isotopic pattern of NADPH relative to NADP^+^ (*eq 1*). The fraction of NADPH coming from different pathways was calculated as described previously^14^:

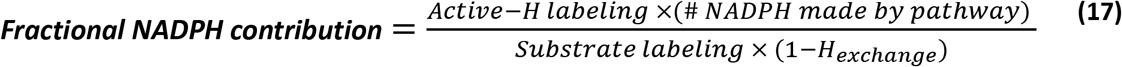

\# NADPH made by pathway is 2 for the oxidative PPP and 1 otherwise. H_exchange_ refers to the fraction of NADPH undergoing H-D exchange with water. Experimental measurements of NADPH and fat labeling after D_2_O infusion gave out *H_exchange_* in liver and brown adipose (*eq 7*; Sup. Fig. 1h). Tissue glucose-6-phosphate labeling was used as substrate labeling for calculating the oxPPP contribution; tissue serine M+2 and M+3 labeling (summed) was used as substrate labeling for calculating the serine contribution. No correction was made for the deuterium kinetic isotope effect.

## Author contributions and information

Z.Z., T.T. and J.D.R. came up with the general approach and tracing strategy. Z.Z and T.T. designed and performed all the *in vivo* and *in vitro* isotope tracing studies and other. Z.Z. conducted most of the sample preparation and data analysis. X.X. performed folate species measurement. X.Z. performed all the acetate measurement. L.Y. contributed to *in vivo* one-carbon metabolism study. G.X., G. J. T. and M. F. C. contributed to *in vitro* tracing experiments. Z.Z., T.T., and J.D.R. wrote the manuscript with help from all authors. Correspondence and requests for materials should be addressed to J.D.R.

## Declaration of interests

J.D.R. is a consultant to Pfizer and an advisor and stock owner in Colorado Research Partners, L.E.A.F. Pharmaceuticals, Rafael Pharmaceuticals, Raze Therapeutics, Kadmon Pharmaceuticals, and Agios Pharmaceuticals. J.D.R. is co-inventor of SHIN2 and related SHMT inhibitors, which have been patented by Princeton University.

## Acknowledgement

This work was supported by NIH Pioneer award 1DP1DK113643 and Pfizer, Inc. T.T. is supported by NIH fellowship 1F32DK118856-01A1. L.Y. was supported by a postdoctoral fellowship from the New Jersey Commission on Cancer Research. We thank Sheng (Tony) Hui, Cholsoon Jang, Li Chen, Yihui Shen, Lin Wang, Caroline Bartman and all other Rabinowitz lab members for advice.

