## Supplementary figures for "Serine catabolism generates NADPH to support hepatic lipogenesis"

### Supplementary Information

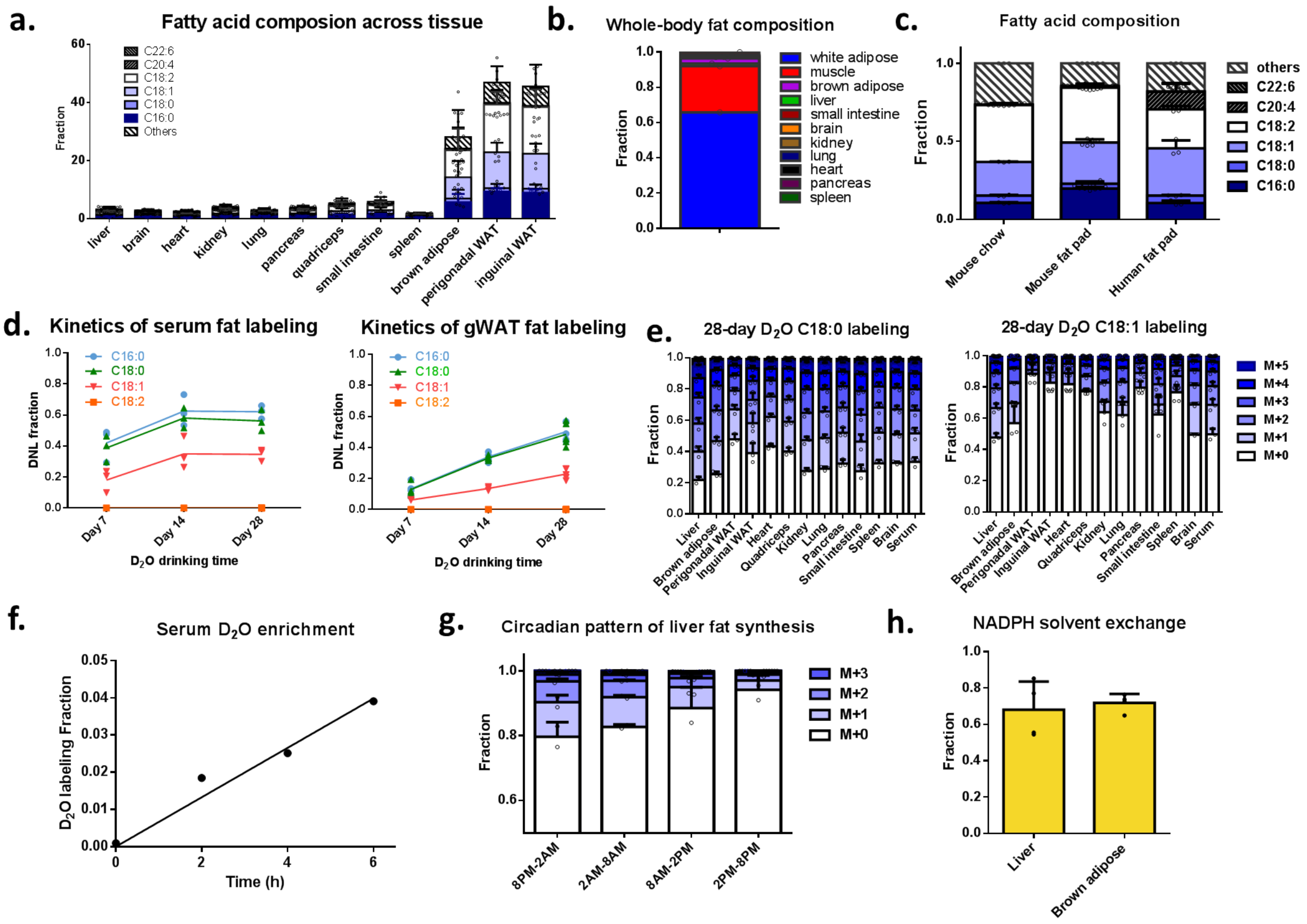


**Supplementary Figure 1. Fatty acid composition and $D_2O$ labeling across tissues and time.**

a. Fraction of tissue weight that is fat, and composition of those fatty acids.
b. Fat is stored in white adipose tissue, and to a lesser extent in muscle. Muscle values are calculated assuming quadriceps is representative of whole-body muscle.
c. Fatty acid composition of mouse chow, mouse inguinal white adipose tissue, and human inguinal white adipose tissue.
d. Fatty acid labeling from $D_2O$ reaches steady-state more quickly in serum than in white adipose, while different fatty acid species reach steady-state at a similar rate.
e. C18:0 and linoleate (C18:1) labeling pattern across tissues after 4 weeks of $D_2O$ drinking
f. Labeling of serum $D_2O$ increases linearly over time for the first 6 h of infusion.
g. Circadian pattern of liver fat synthesis based on 6 h $D_2O$ infusion.
h. NADPH solvent exchange fraction calculated from 12 h $D_2O$ infusion in liver and brown adipose.

Mean ± s.d., n=4 mice for steady-state analysis, composition analysis and NADPH exchange fraction measurement; n=2 mice for 6 h $D_2O$ infusions.

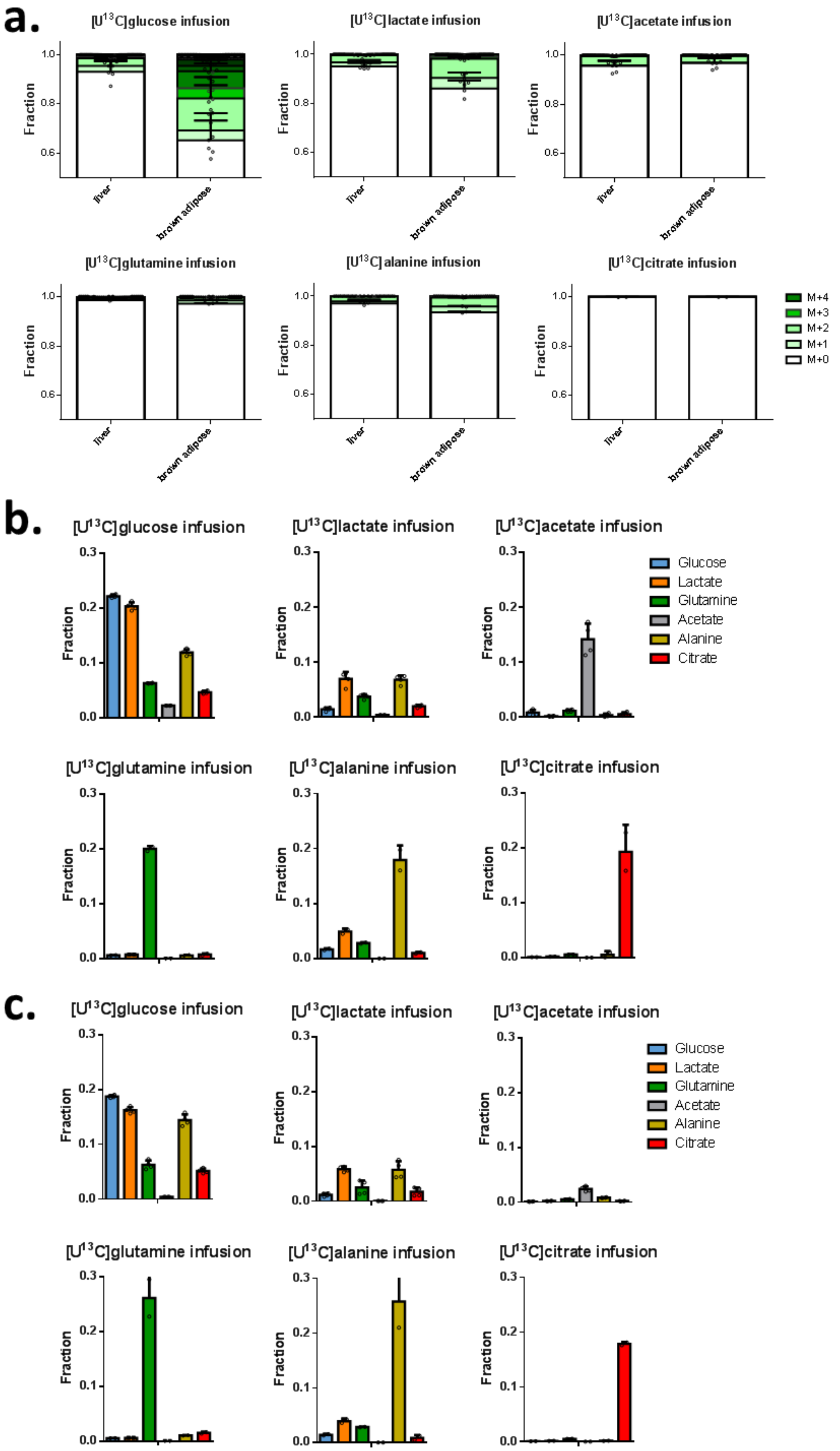
a.
[U13C]glucose infusion
[U13C]lactate infusion
[U13C]acetate infusion
[U13C]glutamine infusion
[U13C]alanine infusion
[U13C]citrate infusion
Fraction
1.0
0.8
0.6
liver
brown adipose
M+4
M+3
M+2
M+1
M+0
b.
[U13C]glucose infusion
[U13C]lactate infusion
[U13C]acetate infusion
[U13C]glutamine infusion
[U13C]alanine infusion
[U13C]citrate infusion
Fraction
0.3
0.2
0.1
0.0
Glucose
Lactate
Glutamine
Acetate
Alanine
Citrate
c.
[U13C]glucose infusion
[U13C]lactate infusion
[U13C]acetate infusion
[U13C]glutamine infusion
[U13C]alanine infusion
[U13C]citrate infusion
Fraction
0.3
0.2
0.1
0.0
Glucose
Lactate
Glutamine
Acetate
Alanine
Citrate

**Supplementary Figure 2. C16:0 and circulating metabolite labeling from different infused carbon tracers.**

a. C16:0 labeling in liver and brown adipose from different $^{13}C$ tracers (each 12 h).

b. Corresponding tail vein serum circulating metabolite labeling.

c. Corresponding portal vein serum circulating metabolite labeling.

All data are mean ± s.d., [U-$^{13}C$]glucose (n=4), lactate (n=4), glutamine (n=2), acetate (n=4), alanine (n=2) and citrate (n=2).

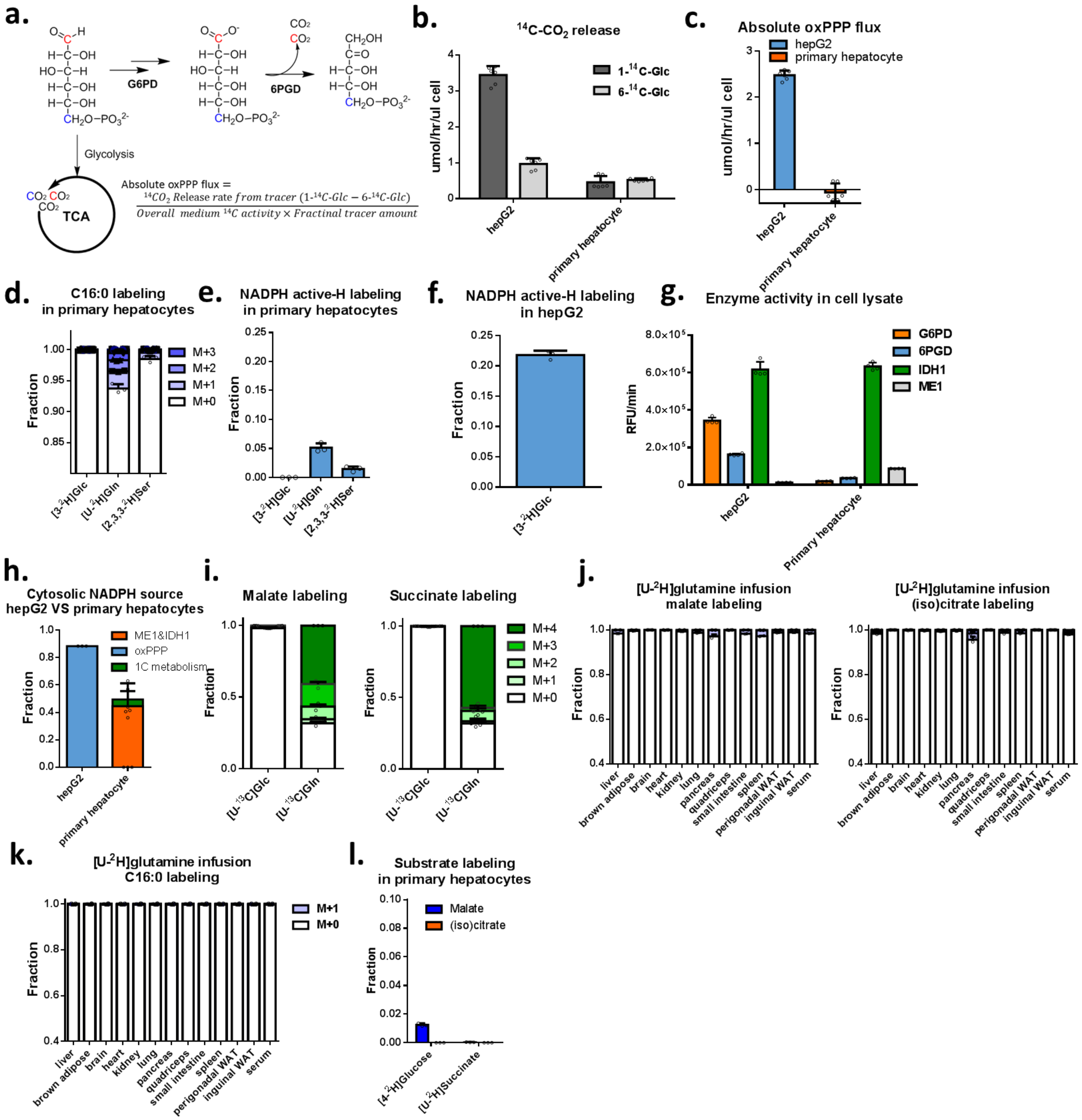


**Supplementary Figure 3. Hepatocytes are deficient in the oxPPP, [U-$^2$H]glutamine tracer works only in cultured hepatocytes but not *in vivo.***

a. Scheme for $^{14}$C experiment to examine absolute oxPPP flux.

b. $^{14}CO_2$ release flux from [1-$^{14}$C]glucose and [6-$^{14}$C]glucose in hepG2 and primary hepatocytes.

c. Absolute oxPPP flux in hepG2 and primary hepatocytes.

d. C16:0 labeling over 6 h from different tracers in primary hepatocytes.

e. NADPH active-H labeling over 2.5h from different tracers in primary hepatocytes.

f. NADPH active-H measurement over 2.5 h from [3-$^2$H]glucose in hepG2 cells.

g. Enzymatic activity of G6PD, 6PGD, IDH1 and ME1 in lysates from hepG2 cells and primary murine hepatocytes.
h. Hydride sources supporting *de novo* lipogenesis in hepG2 cells and primary hepatocytes, correcting for substrate labeling and H-D exchange between NADPH and water.
i. Glutamine is the dominant TCA substrate in cultured primary hepatocytes.
j. Minimal malate and (iso)citrate labeling from [U-$^{2}$H]glutamine infusion across tissues *in vivo*.
k. C16:0 labeling from [U-$^{2}$H]glutamine infusion across tissues.
l. Minimal malate and (iso)citrate labeling from [4-$^{2}$H]glucose and [U-$^{2}$H]succinate in cultured primary hepatocytes.

All data are mean ± s.d., n=2 mice for [U-$^{2}$H]glutamine infusion, 3 replicates for all cell culture work.

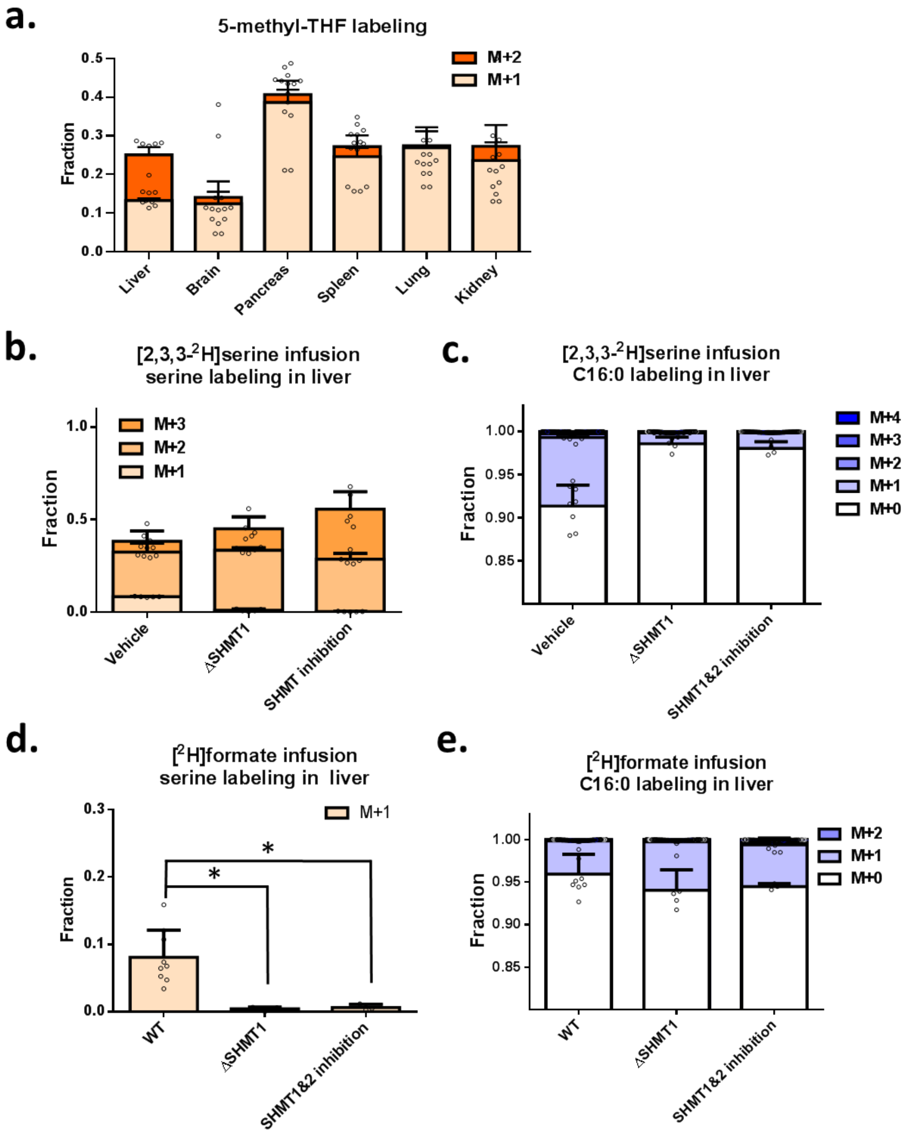


**Supplement Figure 4. 5-methyl-THF, serine, and C16:0 labeling from [2,3,3-$^2$H]serine and [$^2$H]formate**

a. 5-methyl-THF labeling from 12 h [2,3,3-$^2$H]serine. M+2 labeling reflects cytosolic serine catabolism. M+1 labeling reflects the combination of mitochondrial serine catabolism and reversible flux through MTHFD1.
b. Serine labeling in liver from 12 h [2,3,3-$^2$H]serine infusion in in control, whole-body ΔSHMT1 mice, and mice treated with the dual SHMT1/2 inhibitor SHIN2 (3.33 mg/kg/h i.v. infusion).
c. C16:0 labeling in liver from 12 h [2,3,3-$^2$H]serine infusion.
d. As in (b), for [$^2$H]formate infusion.
e. As in (c), for [$^2$H]formate infusion.

Mean± s.d. For serine and formate infusions, N=6 for wild type, N=4 for ΔSHMT1, N=4 for SHIN2 treatment in all experiments. n.s., $P>0.05$; asterisk, $P<0.05$; two asterisks, $P<0.01$; three asterisks, $P<0.005$.
